# Phosphatidylinositol 3,5-Bisphosphate Regulates Yeast Vacuole Fusion at the Transition between *trans*-SNARE Complex Formation and Hemifusion

**DOI:** 10.1101/390062

**Authors:** Gregory E. Miner, Katherine D. Sullivan, Annie Guo, Brandon C. Jones, Matthew L. Starr, Rutilio A. Fratti

## Abstract

Phosphoinositides (PIs) regulate myriad cellular functions including membrane fusion, as exemplified by the yeast vacuole, which uses various PIs at different stages of fusion. In light of this, the effect of phosphatidylinositol 3,5-bisphosphate [PI(3,5)P_2_] on vacuole fusion remains unknown. PI(3,5)P_2_ is made by the PI3P 5-kinase Fab1/PIKfyve and has been characterized as a regulator of vacuole fission during hyperosmotic shock where it interacts with the TRP family Ca^2+^ channel Yvc1. Here we demonstrate that exogenously added dioctanoyl (C8) PI(3,5)P_2_ abolishes homotypic vacuole fusion. This effect was not linked to interactions with Yvc1, as fusion was equally affected using *yvc1Δ* vacuoles. Thus, the effects of C8-PI(3,5)P_2_ on fusion versus fission operate through distinct mechanisms. Further testing showed that C8-PI(3,5)P_2_ inhibited vacuole fusion after the formation of *trans*-SNARE pairs. Although SNARE complex formation was unaffected we found that C8-PI(3,5)P_2_ strongly inhibited hemifusion. Overproduction of endogenous PI(3,5)P_2_ by the *fab1^T2250A^* hyperactive kinase mutant also inhibited at the hemifusion stage, bolstering the model in which PI(3,5)P_2_ inhibits fusion when present elevated levels. Taken together, this work identifies a novel function for PI(3,5)P_2_ as a negative regulator of vacuolar fusion. Moreover, it suggests that this lipid acts as a molecular switch between fission and fusion.

## INTRODUCTION

The ability of lipids, specifically the phosphoinositides (PIs) to regulate membrane homeostasis has been well established. Classically, PI-dependent regulation is associated with the recruitment of soluble proteins to various membranes in order to promote the protein’s function. However, a growing body of work now shows that PIs and other regulatory lipids contribute to a wide array of functions including transcriptional regulation (Carman and Han, 2009), protein sequestering to inhibit function (Starr et al., 2016), microdomain formation (Fratti et al., 2004; Simons and Toomre, 2000), modulating the physical properties of membranes (Andersen and Koeppe, 2007; Zick et al., 2014), and regulating the activity of membrane anchored proteins (Dong et al., 2010; Kiontke et al., 2017; Li et al., 2014). The rapid modifications of lipids by specific kinases, phosphatases, and lipases adds an additional layer of complexity by affecting the spatiotemporal control of cellular events. In the context of *Saccharomyces cerevisiae* vacuolar membrane homeostasis two of the important PI species are PI3P and PI(3,5)P_2_. These lipids appear to play opposing roles as PI3P is characterized as a positive regulatory of fusion (Boeddinghaus et al., 2002; Faber et al., 2009; Fratti et al., 2004; Fratti and Wickner, 2007), whereas PI(3,5)P_2_ promotes vacuolar fission (Bonangelino et al., 2002; Gary et al., 1998).

PI(3,5)P_2_ is found predominately in the late endosome and vacuole where it is made exclusively by the PI3P 5-kinase Fab1 (Gary et al., 1998; Yamamoto et al., 1995). Deletion of *FAB1* results in a complete loss of PI(3,5)P_2_ as well as a drastic enlargement of the vacuole. PI(3,5)P_2_ can be dephosphorylated to PI3P by the PI(3,5)P_2_ 5-phosphatase Fig4 (Gary et al., 2002). Surprisingly, the deletion of *FIG4* leads to an overall decrease in PI(3,5)P_2_ levels (Duex et al., 2006a; Duex et al., 2006b; Efe et al., 2007). This is due to the required presence of Fig4 along with Vac7, Vac14, and Atg18 in the Fab1 complex to yield maximum PI3P 5’-kinase activity (Botelho et al., 2008; Jin et al., 2008; Sbrissa et al., 2007).

The enlarged vacuolar phenotype seen with the loss of Fab1 indicated a role for PI(3,5)P_2_ in membrane fission. Further work showed that hypertonic conditions, which induce vacuolar fission, lead to a rapid increase in PI(3,5)P_2_ levels (Bonangelino et al., 2002; Dove et al., 1997; Duex et al., 2006b). This rise in PI(3,5)P_2_ triggers a release of Ca^2+^ through the vacuolar Ca^2+^ exporter Yvc1 (Dong et al., 2010). Deletion of either Fab1 or Yvc1 attenuates the ability of hypertonic conditions to cause membrane fission, indicating that the increase in PI(3,5)P_2_ and subsequent release of Ca^2+^ are essential for membrane fission. PI(3,5)P_2_ also directly binds to the vacuole specific V-ATPase V_0_ subunit, Vph1 to trigger V-ATPase assembly (Li et al., 2014). Vacuolar acidification itself has been shown to be an essential component of membrane fission as inhibition of V-ATPase H^+^ pumping activity leads to fission defects (Baars et al., 2007).

In this study we examined the ability of PI(3,5)P_2_ to regulate membrane fusion. We found that addition of exogenous (C8) PI(3,5P)_2_ inhibited vacuolar fusion in a dose dependent fashion. This inhibition was independent of Yvc1 as vacuoles lacking the TRP Ca^2+^ channel were equally sensitive to PI(3,5)P_2_. An examination of the distinct stages of membrane fusion indicated that PI(3,5)P_2_ functions after *trans*-SNARE pairing but before hemifusion. Taken together with the role of PI(3,5)P_2_ during osmotic shock, we propose that this lipid acts as a molecular switch which promotes fission while simultaneously inhibiting fusion.

## RESULTS

### Dioctanoyl PI(3,5)P2 inhibits vacuole homotypic fusion

Yeast vacuole fusion is promoted by a group of regulatory lipids including PI3P, PI4P, PI(4,5)P_2_, ergosterol and diacylglycerol, whereas vacuole fission during hyperosmotic shock is regulated by the PI(3,5)P_2_ (Bonangelino et al., 2002; Dove et al., 1997; Duex et al., 2006b; Wickner, 2010). This lipid is made sparingly and turned over rapidly, a likely indication that its prolonged presence or elevated concentration could have a negative impact on vacuole homeostasis. PI(3,5)P_2_ is also made under isotonic conditions, albeit at very low concentrations (Bonangelino et al., 2002; Duex et al., 2006b). Although much is known about PI(3,5)P_2_, its effect during vacuole fusion is unclear. Here we tested the effects of adding exogenous dioctanoyl (C8) PI(3,5)P_2_ to *in vitro* vacuole homotypic fusion reactions. We found that C8-PI(3,5)P_2_ inhibited vacuole fusion in a dose dependent manner with an IC_50_ of 110.2 µM (Fig. 1A, black circles). During hyperosmotic shock, PI(3,5)P_2_ is known to activate the TRP Ca^2+^ channel Yvc1. To test whether the inhibitory effect of PI(3,5)P_2_ on vacuole fusion was due to Yvc1 activation, we used vacuoles from *yvc1Δ* yeast. The addition of C8-PI(3,5)P_2_ to *yvc1Δ* vacuoles had a nearly identical inhibitory effect on fusion relative to wild type vacuoles. The IC_50_ of C8-PI(3,5)P_2_ on *yvc1Δ* vacuoles was 126.2 µM (Fig. 1A, red squares). Thus, we can conclude that the effect of C8-PI(3,5)P_2_ is independent of activating Yvc1 function, *i.e.* enhanced Ca^2+^ efflux. As a control we used the PI(3,5)P_2_ precursor PI3P. Wild type vacuoles were treated with a dose curve of C8-PI3P and tested for fusion efficiency. Unlike the inhibitory effects of PI(3,5)P_2_, there was no significant reduction in fusion efficiency when C8-PI3P was added (Fig. 1B). This suggests that the effects of PI(3,5)P_2_ on fusion were specific to the lipid head group.

**Figure 1.**
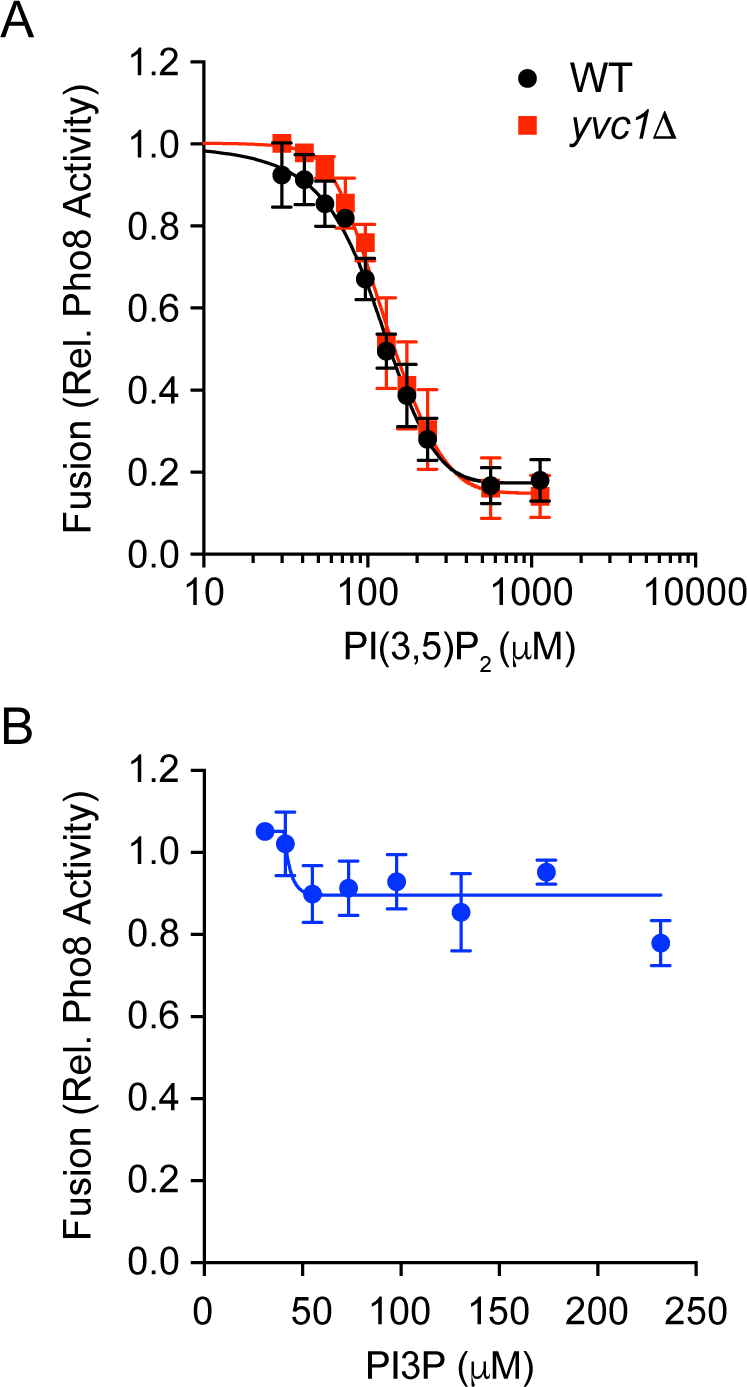
Short chain PI(3,5)P_2_ inhibits vacuole homotypic fusion. Vacuoles isolated from WT or *yvc1Δ* fusion reporter backgrounds were incubated with buffer alone or a dose curve of C8-PI(3,5)P_2_ (A) or C8-PI3P (B) as indicated. Fusion reactions were incubated for 90 min at 27°C. After incubation membranes were solubilized and incubated with *p*-nitrophenyl phosphate to measure Pho8 activity. *p-*nitrophenolate was measured at OD_400_. Error bars are S.E.M. (n=3).

### C8-PI(3,5)P_2_ inhibits vacuole fusion at the docking stage

In a previous study we found that adding short chain phosphatidic acid (PA) inhibited vacuole fusion at the priming stage by preventing the recruitment of Sec18 to *cis*-SNARE complexes (Starr et al., 2016). To begin to determine at which stage PI(3,5)P_2_ inhibited fusion, we first tested SNARE priming. When Sec18 is recruited to inactive SNAREs for priming, it associates with *cis*-SNARE complexes though its interactions with the adaptor protein Sec17/α-SNAP. Once Sec18 hydrolyses ATP to dissociate SNAREs, the bridging Sec17 molecules are released from the membrane (Mayer et al., 1996). Thus, priming can be measured by the loss of Sec17 from the membrane. Here, fusion reactions were incubated with reaction buffer, 232 µM C8-PI(3,5)P_2_ or 1 mM *N*-ethylmaleimide (NEM), a known inhibitor of priming (Paumet et al., 2000; Starr et al., 2016). Individual reactions were incubated for specific times after which they were centrifuged to separate the membrane bound (pellet) and solubilized (supernatant) fractions of Sec17 that were then detected by Western blotting. These experiments showed that PI(3,5)P_2_ did not negatively affect SNARE priming, as similar levels of Sec17 were released relative to the buffer control (Fig. 2A-B). By contrast, SNARE priming was inhibited by NEM as shown by the complete block of Sec17.

**Figure 2.**
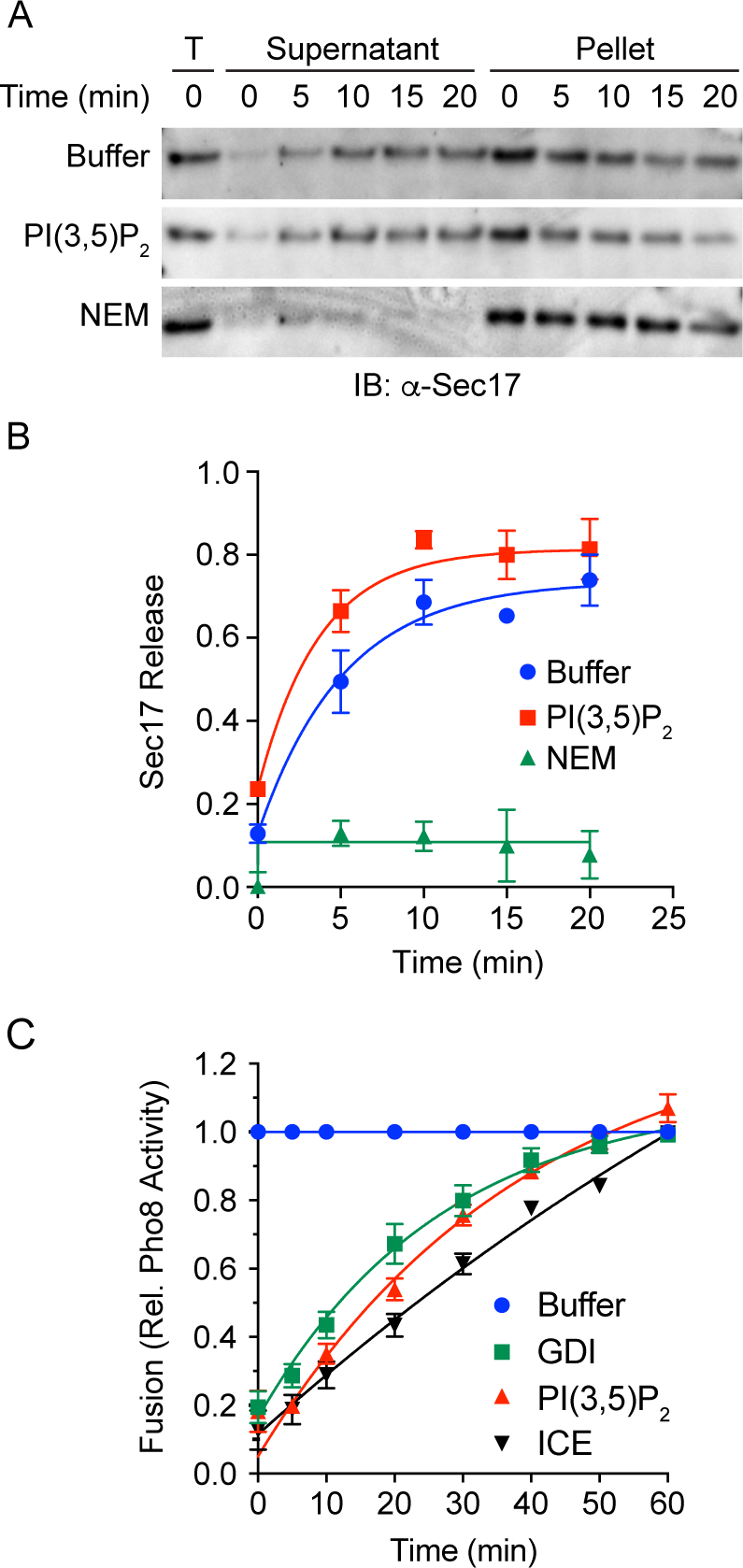
C8-PI(3,5)P_2_ inhibits vacuole fusion at the docking stage. (A) Vacuoles from BJ3505 were monitored for the release of Sec17 from the membrane upon SNARE priming. Fusion reactions containing 3 µg of vacuoles (by protein) were incubated with reaction buffer, 232 µM C8-PI(3,5)P_2_ or 1 mM NEM. Vacuoles were incubated at 27°C for the indicated times after witch the organelles were pelleted by centrifugation and released proteins in the supernatant were separated from the membrane bound fraction. The membrane pellets were resuspended in volumes of reaction buffer equal to the supernatant. Both fractions were mixed with SDS loading buffer and resolved by SDS-PAGE. Sec17 was detected by Western blotting and the amount released was calculated by densitometry. (B) Normalized values were averaged and plotted over time of incubation. (C) Gain of resistance kinetic vacuole fusion assays were performed in the presence of reaction buffer, 2 µM GDI, or 232 µM C8-PI(3,5)P_2_. Reactions were incubated at 27°C or on ice for 90 min. Reagents were added at the indicated time points. Fusion inhibition was normalized to the reactions receiving buffer alone. Data were fit with first order exponential decay. Error bars are S.E.M. (n=3).

To further resolve at which stage of the pathway PI(3,5)P_2_ inhibited fusion we performed a temporal gain of resistance assay (Sasser et al., 2012a). Here, inhibitors were added to individual reactions at different time-points throughout the incubation period. Fusion reactions become resistant to a reagent once the target of the inhibitor carried out its function. Thus, inhibitors of early events (*e.g.* priming) would lose their effects before inhibitors of later stages (*e.g.* hemifusion). Through this approach we found that PI(3,5)P_2_ inhibited late in the docking stage, as its resistance curve lied between that of the early docking inhibitor GDI and the ice curve (Fig. 2C). GDI extracts the Rab Ypt7 during fusion to prevent the early stages of docking/tethering and prevents the formation of *trans*-SNARE pairs between vacuoles (Eitzen et al., 2000; Eitzen et al., 2001; Haas et al., 1995). The uninhibited control was normalized to 1 for each time point. These data indicate that PI(3,5)P_2_ inhibits vacuole fusion after Ypt7-dependent vacuole association.

### C8-PI(3,5)P_2_ does not inhibit *trans*-SNARE complex formation

Due to the indication that PI(3,5)P_2_ inhibited vacuole fusion after tethering we next examined if SNARE complex formation occurred. For this, we used our established GST-Vam7 bypass assay where fusion reactions are blocked at the priming stage with anti-Sec17 IgG (Fratti and Wickner, 2007; Fratti et al., 2007; Karunakaran and Fratti, 2013; Miner et al., 2016). The addition of GST-Vam7 allows for the formation of new *trans* complexes with free SNAREs that are not in *cis*. Between the addition of anti-Sec17 and GST-Vam7, we further treated individual reactions with GDI to block tethering or PI(3,5)P_2_. Once GST-Vam7 was added, the reactions were incubated for 60 min at 27°C or on ice as indicated. After incubation, the reactions were placed on ice and solubilized as described in the materials and methods section. GST-Vam7 SNARE complexes were isolated with reduced glutathione beads and proteins were resolved by SDS-PAGE. Specific proteins were detected by Western blotting. Figure 3A shows that GST-Vam7 formed complexes with its cognate SNAREs Vam3 and Nyv1 when incubated at 27°C (lane 3), whereas incubation on ice prevented complex formation as seen previously (Fratti and Wickner, 2007; Fratti et al., 2007). While GDI inhibited SNARE complex formation the addition of PI(3,5)P_2_ had no effect on the interactions of these proteins (lane 4 vs 5). Quantitation of multiple experiments showed that PI(3,5)P_2_ indeed had no significant effect on SNARE pair formation (Fig. 3B).

**Figure 3.**
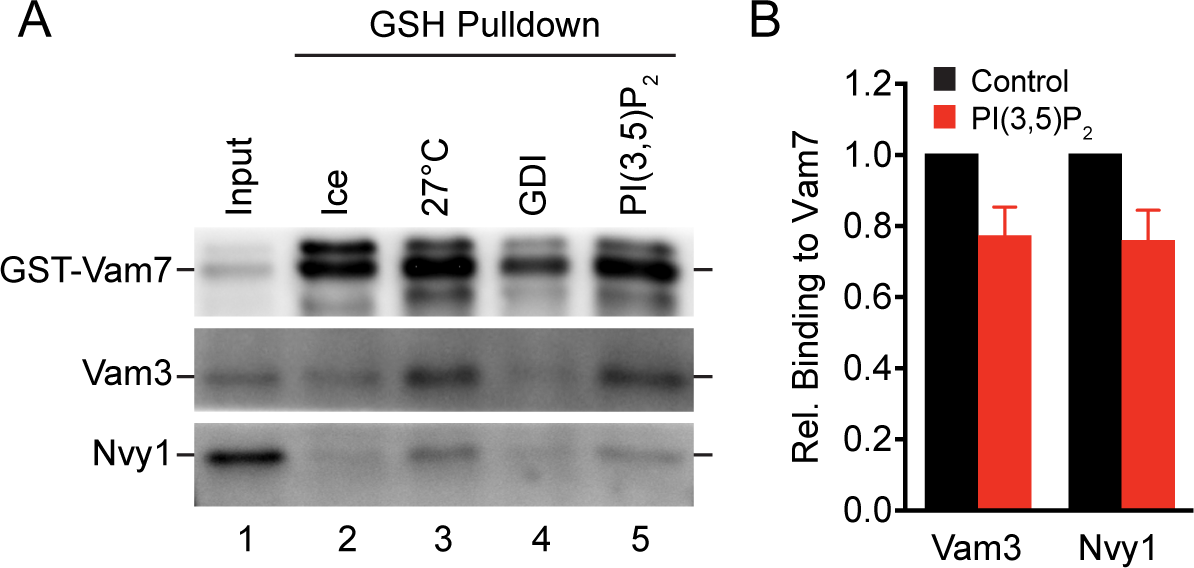
PI(3,5)P_2_ does not inhibit *trans*-SNARE complex formation. (A) Large scale vacuole fusion reactions (6x) were incubated with anti-Sec17 IgG to block SNARE priming. After incubating for 15 min at 27°C, select reactions were further treated with either reaction buffer, 2 µM GDI or 232 µM C8-PI(3,5)P_2_ and incubated for an additional 70 min. One reaction remained on ice for the duration of the assay. Reactions were then processed for glutathione pulldown of GST-Vam7 protein complexes. Isolated protein complexes were resolved by SDS-PAGE and probed for the presence of the SNAREs Vam3 and Nyv1. (B) Quantitation of SNARE complex formation in the presence or absence of C8-PI(3,5)P_2_. Error bars are S.E.M. (n=3).

### Hemifusion is blocked by C8-PI(3,5)P_2_

Previous studies have shown that lysophosphatidylcholine (LPC) blocked fusion late in the pathway, with a gain of resistance curve similar to what we saw with PI(3,5)P_2_ (Reese and Mayer, 2005). While LPC permitted SNARE complex assemble, Reese and Mayer showed that LPC blocked hemifusion in a dose dependent manner. We next examined if PI(3,5)P_2_ had a similar effect to LPC in blocking hemifusion after allowing *trans*-SNARE complex formation. Hemifusion was measured by the dequenching of Rhodamine labeled-phosphatidylethanolamine (Rh-PE) on the outer leaflets of vacuoles. Rh-PE is loaded onto the outer leaflet of vacuoles at a concentration that causes the dye to self-quench. Rh-PE labeled vacuoles were incubated with an 8-fold excess of unlabeled vacuoles. Upon the fusion of the outer leaflets, the Rh-PE was diluted to restore fluorescence. Our results showed that PI(3,5)P_2_ potently blocked hemifusion similar to what was seen with LPC (Fig. 4A-B). As a negative control we used GDI to block upstream of hemifusion at the tethering stage. These data indicate that PI(3,5)P_2_ blocks fusion at a stage after *trans*-SNARE pairing but before lipid mixing can occur.

**Figure 4.**
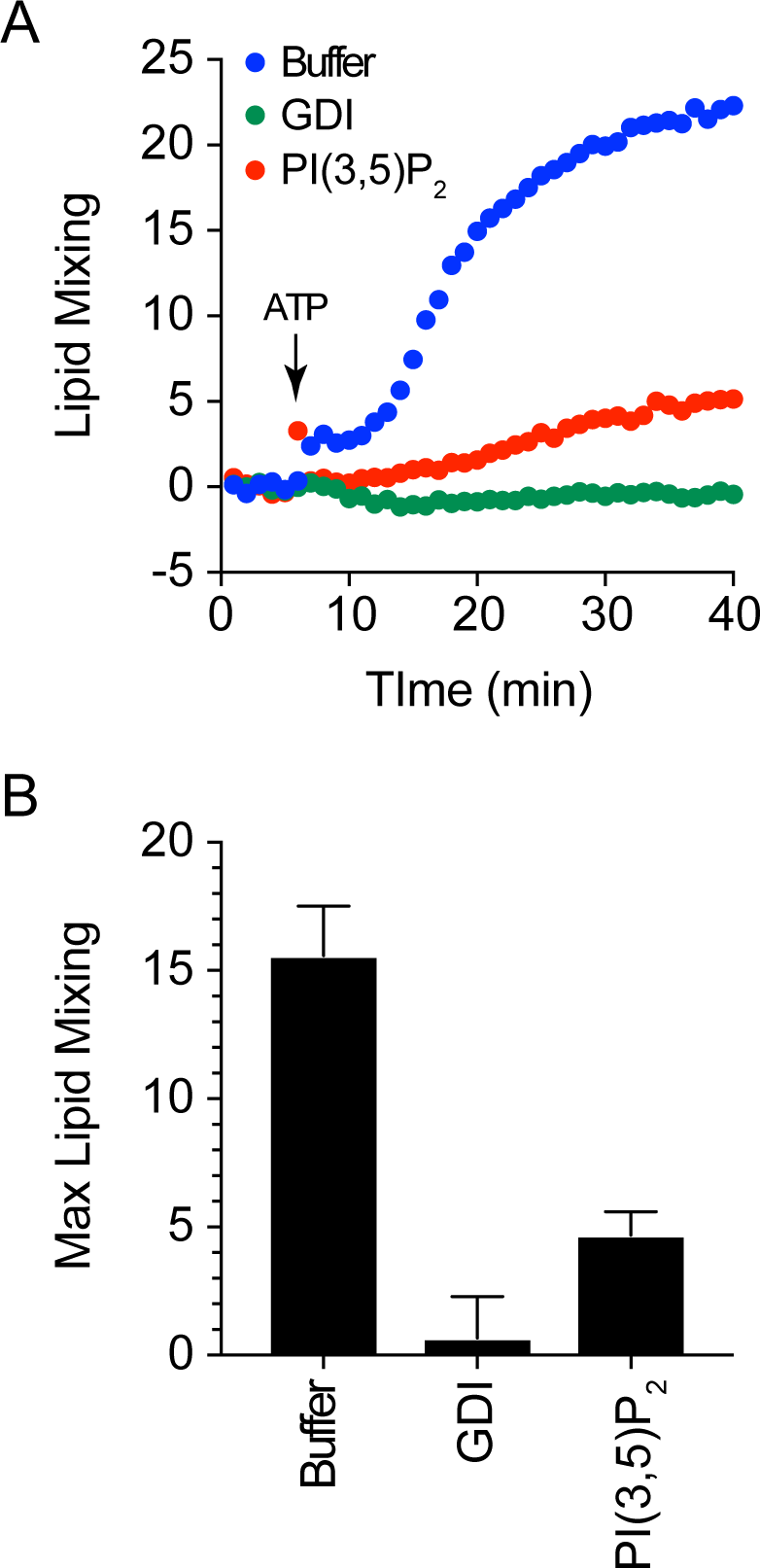
Hemifusion is blocked by C8-PI(3,5)P_2_. (A) Lipid mixing assay measuring outer leaflet fusion, *i.e.* hemifusion, were performed using vacuoles labeled with Rh-PE. Labeled vacuoles were incubated with an 8-fold excess of unlabeled vacuoles. Dilution of Rh-PE upon lipid mixing leading to fluorescence dequenching was measured using a plate reader. Lipid mixing reactions were treated with either reaction buffer, 2 µM GDI or 232 µM C8-PI(3,5)P_2_ and incubated for 40 min. Reactions were started with the addition of ATP after 5 min. Fluorescence was measured every 60 seconds and plotted against time. (B) Quantitation of average maximum Rh-PE fluorescence. Error bars are S.E.M. (n=3).

### Fab1 Mutants Affect Hemifusion

To verify that an excess of PI(3,5)P_2_ inhibits hemifusion, we used vacuoles harvested from yeast expressing Fab1 with the hyperactive kinase mutation T2250A (Lang et al., 2017). As predicted, vacuoles expressing *fab1^T2250A^* failed to undergo lipid mixing (Fig. 5A-B). This is attributed to the increased production of PI(3,5)P_2_ and is consistent with our data using exogenous lipid. We next tested whether the lack of PI(3,5)P_2_ would have the opposite effect on lipid mixing. To address this, we used vacuoles from yeast that expressed the kinase dead mutant *fab1^EEE^* (Li et al., 2014). We found that *fab1^EEE^* vacuoles were also blocked for lipid mixing (Fig. 5C-D). This suggests that some PI(3,5)P_2_ production is needed for fusion, but an excess is inhibitory. These data illustrate that PI(3,5)P_2_ levels have a direct effect on the ability of vacuoles to transition through the hemifusion stage toward full content mixing. Finally, we used vacuoles from *fig4Δ* yeast. Cells lacking Fig4 produce reduced amounts of PI(3,5)P_2_ relative to wild type cells due attenuated Fab1 activity in the absence of the phosphatase. This showed that *fig4Δ* vacuoles had reduced lipid mixing (Fig. 5E-F). Together, these data suggest that the hemifusion stage requires some Fab1 activity, but augmented PI(3,5)P2 production is inhibitory.

**Figure 5.**
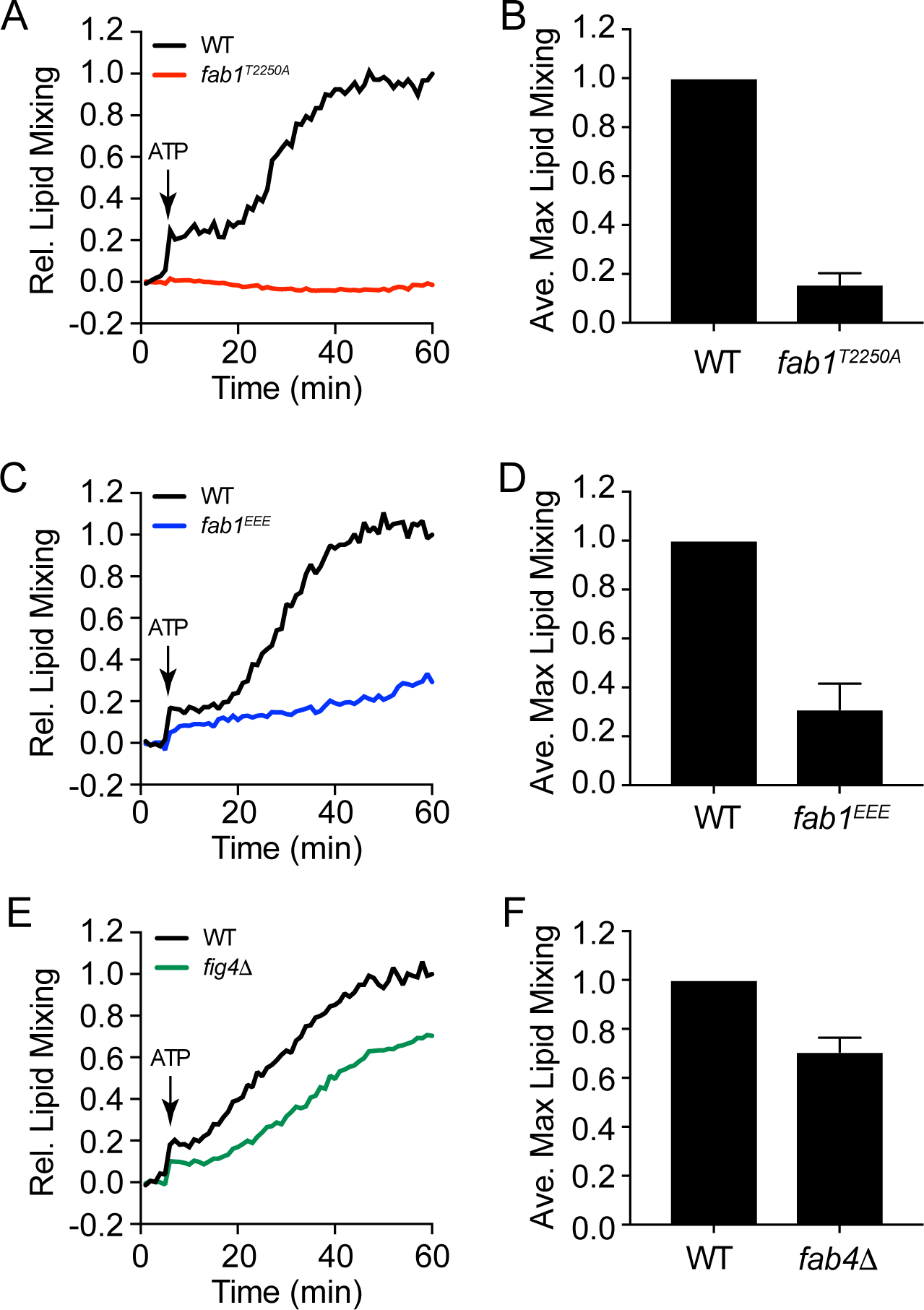
Fab1 mutants block hemifusion. Lipid mixing assays were performed with wild type vacuoles or those harboring the hyperactive kinase mutant *fab1^T2250A^* (A-B), the kinase dead mutant *fab1^EEE^* (C-D), or from *fig4Δ* yeast. Labeled vacuoles were incubated with an 8-fold excess of unlabeled vacuoles as in Figure 4. Lipid mixing reactions were incubated for 60 min and fluorescence was measured every 60 seconds. Reactions were started with the addition of ATP after 5 min. (B, D, F) Quantitation of average maximum Rh-PE fluorescence. Error bars are S.E.M. (n=3).

## DISCUSSION

Regulatory lipids affect many aspects of membrane trafficking, including yeast vacuole homotypic fusion. The fusion of these organelles requires PI3P, PI4P, PI(4,5)P_2_, DAG and ergosterol to establish the vertex microdomain, binding Mon1-Ccz1 for nucleotide exchange activity on Ypt7, Vam7 binding, HOPS association, actin remodeling and promoting SNARE priming (Boeddinghaus et al., 2002; Cabrera et al., 2014; Fratti et al., 2004; Jun et al., 2004; Karunakaran et al., 2012; Karunakaran and Fratti, 2013; Kato and Wickner, 2001; Lawrence et al., 2014; Mayer et al., 2000; Stroupe et al., 2006). For the most part, the effects of regulatory lipids have been found to promote vesicular fusion. An exception to this is the inhibitory role that PA exerts on Sec18 to sequester the priming machinery away from *cis*-SNARE complexes (Starr et al., 2016). In this work we report that vacuole fusion is blocked by PI(3,5)P_2_. Unlike PA, PI(3,5)P_2_ had no effect on SNARE priming. Moreover, SNARE complex formation at the docking stage was unaffected. Our data showed that PI(3,5)P_2_ blocked the fusion after the formation of SNARES and before triggering hemifusion.

How does PI(3,5)P_2_ directly inhibit fusion? One possible mechanism for the inhibition of fusion could be linked to the vacuolar ATPase. Others have reported that PI(3,5)P_2_ stabilizes the interactions of the V1 and V0 components of the V-ATPase (Li et al., 2014). This occurs through the direct interactions of the V0 subunit Vph1 and PI(3,5)P_2_, which promotes binding the V1 complex. The increased assembly of the V1-V0 holoenzyme would in turn deplete free V0 complexes on the vacuole membrane. The significance of altering the levels of free V0 complexes and vacuole fusion lies in the ability of V0 complexes to interact in trans between apposed membranes. Mayer and colleagues originally found that the formation of trans V0-V0 complexes play a role in membrane fusion (Peters et al., 2001). Since their discovery, several other groups using various systems found that V0 plays important non-canonical roles in membrane fusion including the release of insulin (Sun-Wada et al., 2006), phagosome-lysosome fusion (Peri and Nusslein-Volhard, 2008), and the release of exosomes through the fusion of multivesicular bodies with the plasma membrane (Liégeois et al., 2006). The c-rings of the V0 also interact in trans between membranes to form conductance channels through their proteolipid pores (Couoh-Cardel et al., 2016). Although PI(3,5)P_2_ has not been shown to directly reduce V0-V0 trans complexes, it stands to reason that the increase in V1-V0 complexes would in turn diminish the former and potentially alter fusion efficiency.

Another potential target for PI(3,5)P_2_ could involve altering Ca^2+^ transport across the vacuole membrane. During salt shock, PI(3,5)P_2_ is rapidly synthesized and interacts with the TRP channel Yvc1 to release vacuolar Ca^2+^ stores (Dong et al., 2010). Because vacuolar Ca^2+^ stores are also released in response to the formation of *trans*-SNARE pairs (Merz and Wickner, 2004), we initially hypothesized that PI(3,5)P_2_ could affect fusion through interactions with Yvc1. However, vacuoles from *yvc1Δ* yeast were equally susceptible to PI(3,5)P_2_ relative to the wild type parents, suggesting that PI(3,5)P_2_ has a different target for the inhibition of fusion. Taken together with the studies on PI(3,5)P_2_ and vacuole fission, we propose that this lipid performs two opposing functions as a molecular switch to go between fission and fusion.

## MATERIALS AND METHODS

### Reagents

Soluble reagents were dissolved in PIPES-Sorbitol (PS) buffer (20 mM PIPES-KOH, pH 6.8, 200 mM sorbitol) with 125 mM KCl unless indicated otherwise. Anti-Sec17 IgG (Mayer et al., 1996), and Pbi2 (Slusarewicz et al., 1997) were prepared as described previously. C8-PI3P (1,2-dioctanoyl-phosphatidylinositol 3-phosphate), and C8-PI(3,5)P_2_ (1,2-dioctanoyl-phosphatidylinositol 3,5-bisphosphate) were purchased from Echelon Inc. Recombinant GST-Vam7 and GDI were prepared as previously described (Fratti and Wickner, 2007; Fratti et al., 2007; Starai et al., 2007). Lissamine rhodamine (Rh-PE) was from ThermoFisher. NEM (N-ethylmaleimide) was from Sigma.

### Strains

BJ3505 (Jones et al., 1982) and DKY6281 (Haas et al., 1994) were used for fusion assays (Table 1)*. YVC1* was deleted by homologous recombination using PCR products amplified from pFA6-KanMX6 with primers 5’-YVC1-KO (5’-ATTCAGTTATAAAATATAATATTACTAGAACAGGAGCATTCGGATCCCCGGGTTAATTAA–3’) and 3’-YVC1-KO (5’-TTCTGAGAAATTAATTAAGCAGTATTTGAACACATGTCGTGAATTCGAGCTCGTTTAAAC–3’) with homology flanking the *YVC1* coding sequence. The PCR product was transformed into BJ3505 and DKY6281 yeast by standard lithium acetate methods and plated on YPD media containing G418 (250 µg/ml) to generate BJ3505 *yvc1Δ::kanMX6* (RFY74) and DKY6281 *yvc1Δ::kanMX6* (RFY75). Similarly, *FAB1* was deleted by recombination using the primers 5’-FAB1-KO (5’-AGGTAGCTTCCATCCTGTACATGCAAGACCGTCACACAGCCGGATCCCCGGGTTAATTAA–3’) and 3’-FAB1-KO (5’-TAAAAAAAAGTTACAGAATATAACTTGTACACGTTTATGTGAATTCGAGCTCGTTTAAAC–3’) to generate BJ3505 *fab1Δ::kanMX6* (RFY76). RFY76 was transformed with a plasmid encoding the hyperactive kinase mutant *fab1T2250A* (pRS416-FAB1-T2250A) and grown in selective media lacking uracil to make RFY78 (Lang et al., 2017). Similarly, a plasmid encoding the kinase dead mutant *fab1^EEE^* (pRS416-FAB1-EEE) was transformed into RFY76 to make RFY80 (Li et al., 2014). Plasmids harboring Fab1 mutants were a gift from Dr. Lois Weisman (University of Michigan). *FIG4* was deleted through recombination as described above using PCR products amplified from pAG32 with the primers 5’-FIG4-KO (5’-AACGTAGTAAC GTAACGCAAAGCAAAAAAGAAACAGGACACGGATCCCCGGGTTAATTAA-3’) and 3’-FIG4-KO (TTGACCGAA-TATTT AAATTTCTATTTAAGAGTCATATAAAGAATTCGAGCTCGTTTAAAC) to generate BJ3505 *fig4Δ::kanMX6* (RFY82). Transformants were grown in YPD with hygromycin (250 µg/ml).

**Table 1.**
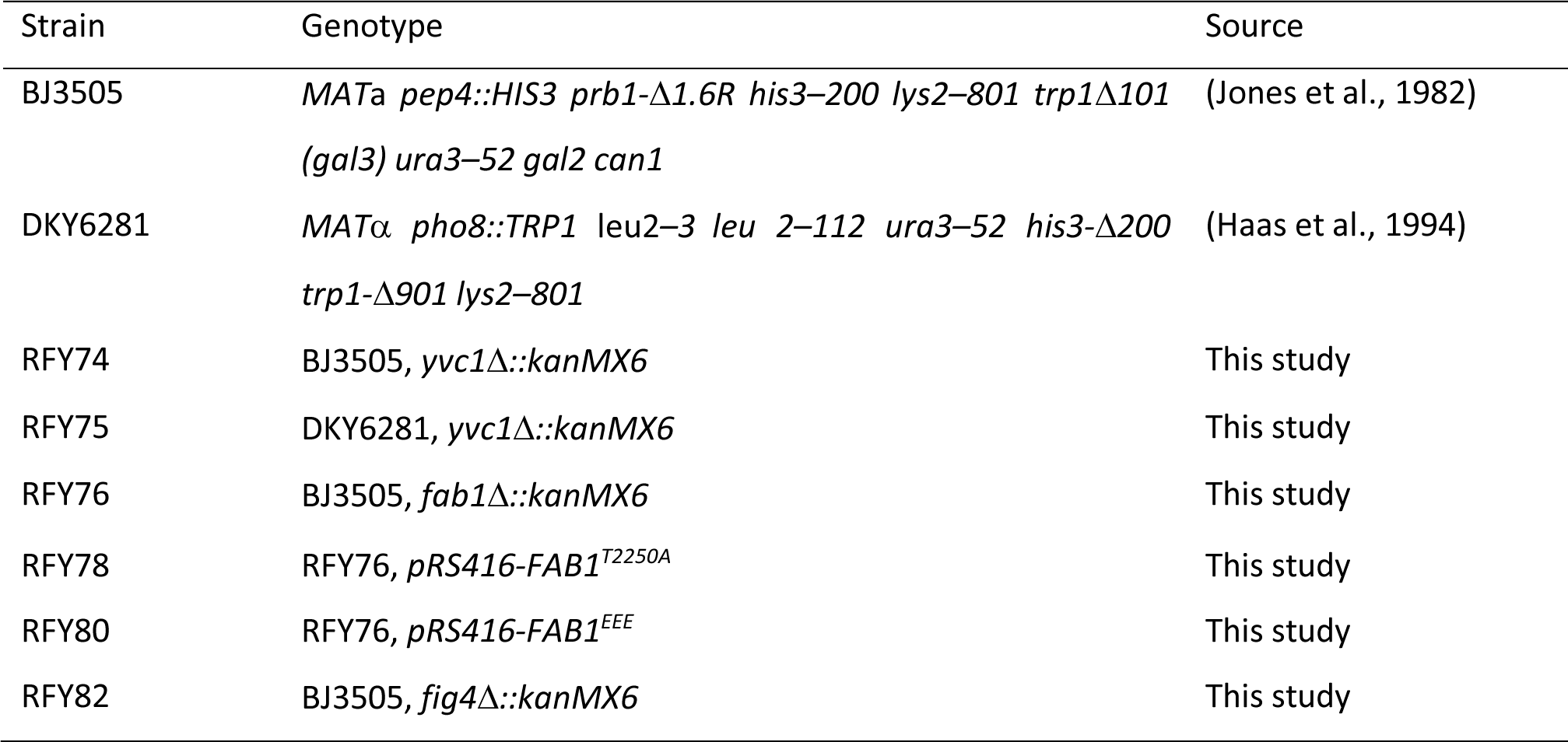
Yeast strains used in this study

### Vacuole Isolation and In-vitro fusion assay

Vacuoles were isolated as described (Haas et al., 1994). *In vitro* fusion reactions (30 µl) contained 3 µg each of vacuoles from BJ3505 (*PHO8 pep4Δ*) and DKY6281 (*pho8Δ PEP4*) backgrounds, reaction buffer 20 mM PIPES-KOH pH 6.8, 200 mM sorbitol, 125 mM KCl, 5 mM MgCl_2_), ATP regenerating system (1 mM ATP, 0.1 mg/ml creatine kinase, 29 mM creatine phosphate), 10 µM CoA, and 283 nM Pbi2 (Protease B inhibitor). Fusion was determined by the processing of pro-Pho8 (alkaline phosphatase) from BJ3505 by the Pep4 protease from DK6281. Fusion reactions were incubated at 27°C for 90 min and Pho8 activity was measured in 250 mM Tris-HCl pH 8.5, 0.4% Triton X-100, 10 mM MgCl_2_, and 1 mM *p*-nitrophenyl phosphate. Pho8 activity was stopped after 5 min by addition of 1 M glycine pH 11 and fusion units were measured by determining the *p-*nitrophenolate produced by detecting absorbance at 400 nm.

### GST-Vam7 SNARE complex isolation

SNARE complex isolation was performed as described previously using GST-Vam7 (Fratti and Wickner, 2007; Fratti et al., 2007; Miner et al., 2016; Miner et al., 2017). Briefly, 5X fusion reactions were incubated with 85 µg/ml anti-Sec17 IgG to block priming. After 15 min, 2 µM GDI or 232 µM C8-PI(3,5)P_2_ was added to selected reactions and incubated for an additional 5 min before adding 150 nM GST-Vam7. After a total of 90 min, reactions were sedimented (11,000 *g*, 10 min, 4°C), and the supernatants were discarded before extracting vacuoles with solubilization buffer (SB: 20 mM HEPES-KOH, pH 7.4, 100 mM NaCl, 2 mM EDTA, 20% glycerol, 0.5% Triton X-100, 1 mM DTT) with protease inhibitors (1 mM PMSF, 10 µM Pefabloc-SC, 5 µM pepstatin A, and 1 µM leupeptin). Vacuole pellets were overlaid with 100 µl SB and resuspended gently. An additional 100 µl SB was added, gently mixed, and incubated on ice for 20 min. Insoluble debris was sedimented (16,000 *g*, 10 min, 4°C) and 176 µl of supernatants were removed and placed in chilled tubes. Next, 16 µl was removed from each reaction as 10% total samples, mixed with 8 µl of 3X SDS loading buffer and heated (95°C, 5 min). Equilibrated glutathione beads (30 µl) were incubated with the remaining extracts (15 h, 4°C, nutation). Beads were sedimented and washed 5X with 1 ml SB (735 *g*, 2 min, 4°C), and bound material was eluted with 40 µl 1X SDS loading buffer. Protein complexes were examined by Western blotting.

### Lipid Mixing

Lipid Mixing assays were conducted using rhodamine B DHPE (Rh-PE; Thermo-Fisher) as described (Karunakaran and Fratti, 2013; Miner et al., 2016; Miner et al., 2017; Sasser et al., 2012a; Sasser et al., 2012b; Sasser et al., 2013). BJ3505 vacuoles (300 µg) were isolated and then incubated in 400 µl of PS buffer containing 150 µM Rh-PE (10 min, 4°C, nutating). Next, 800 µl of 15% Ficoll was added and then transferred to an 11 x 60 mm ultracentrifuge tube, overlaid with 1.2 ml of 8% and 4%, and 0.5 ml of PS buffer. Labeled vacuoles were isolated by centrifugation (105,200 X *g*, 25 min, 4°C, SW-60 Ti rotor) and recovered from the 0-4% Ficoll interface. Lipid mixing assays (90 µl) contained 2 µg of labeled vacuoles and 16 µg of unlabeled vacuoles in fusion buffer. Reaction mixtures were transferred to a black, half volume 96-well flat-bottom microtiter plate (Corning 3686) on ice. The plate was transferred to a fluorescence plate reader at 27°C to start the reactions. Measurements were taken every 60 sec for 40 min yielding fluorescence values (λ_ex_=544 nm; λ_em_=590 nm) at the onset (F_0_) and during the reaction (F_t_). After 40 min 0.45% (vol/vol) Triton X-100 was added and the final 10 measurements were averaged to give the value of fluorescence after infinite dilution (F_TX100_). The relative fluorescence change ΔF_t_/F_TX100_ = (F_t_ - F_0_)/F_TX100_ - F_0_ was calculated.

## ACKNOWLEDGMENTS

We thank Lois Weisman (University of Michigan) for Fab1 plasmids. This research was supported by a grant from the National Institutes of Health (R01-GM101132) and National Science Foundation (MCB 18-18310) to RAF.

